# Systematic and quantitative view of the antiviral arsenal of prokaryotes

**DOI:** 10.1101/2021.09.02.458658

**Authors:** Florian Tesson, Alexandre Hervé, Marie Touchon, Camille d’Humières, Jean Cury, Aude Bernheim

## Abstract

Facing the abundance and diversity of phages, bacteria have developed multiple anti-phage mechanisms. In the past three years, the number of known anti-phage mechanisms has been expanded by at least 5-fold rendering our view of prokaryotic immunity obsolete. Most anti-phage systems have been studied as standalone mechanisms, however many examples demonstrate strains encode not one but several anti-viral mechanisms. How these different systems integrate into an anti-viral arsenal at the strain level remains to be elucidated. Much could be learned from establishing fundamental description of features such as the number and diversity of anti-phage systems encoded in a given genome. To address this question, we developed DefenseFinder, a tool that automatically detects known anti-phage systems in prokaryotic genomes. We applied DefenseFinder to >20 000 fully sequenced genomes, generating a systematic and quantitative view of the anti-viral arsenal of prokaryotes. We show prokaryotic genomes encode on average five anti-phage systems from three different families of systems. This number varies drastically from one strain to another and is influenced by the genome size and the number of prophages encoded. Distributions of different systems are also very heterogenous with some systems being enriched in prophages and in specific clades. Finally, we provide a detailed comparison of the anti-viral arsenal of 15 common bacterial species, revealing drastic differences in anti-viral strategies. Overall, our work provides a free and open-source software, available as a command line tool or, on a webserver. It allows the rapid detection of anti-phage systems, enables a comprehensive description of the anti-viral arsenal of prokaryotes and paves the way for large scale genomics study in the field of anti-phage defense.

## Introduction

Facing the abundance and diversity of phages, bacteria have developed multiple anti-phage mechanisms. Up to 2018, only a few systems were described, including CRISPR-Cas system, Restriction-Modification (RM) and Abortive infection (Abi). In 2018, a landmark study marked the beginning of a new era of discovery by revealing the existence of ten novel anti-phage defense systems^1^. Since then, dozens of novels systems have been unearthed. A majority of these systems were uncovered through the “defense islands” method, using a guilt by association approach^1–8^. Others were discovered individually^9–11^ or by looking into hotspots encoded in mobile genetic element^12–14^. It is thus now recognized that prokaryotic immunity is much more complex than previously perceived with evidence for intracellular signaling regulating defense^5,15^, chemical defense^7,16^, nucleotide depletion^8^, RNA mutations^4^, guardian systems^6^ and the discovery of many prokaryotic defense systems which mechanisms are still unknown^1,4,12^

The discovery of a stockpile of novel anti-phage systems questions our view of how prokaryotes defend themselves against viruses. While many families of systems have been investigated mechanistically, much remains to be uncovered about the anti-viral arsenal at the level of a strain, a species or all prokaryotes. Describing which systems genomes encode will be essential for understanding phage bacteria interactions, in a natural context.

Establishing a holistic, genome centric view of the whole anti-viral arsenal of prokaryotes is currently challenging. Most studies tackling the distribution of defense systems focused on one or a few families of systems. They provide important number regarding their abundance in microbial genome. For example, in a dataset of 38 167 genomes, 4 894 encoded CBASS (13%) and 4446 retrons (11%) ^6,17^. Systems described in ^1^ were found at frequencies ranging from 1.8% (Kiwa) to 8.5% (Gabija)^1^ while RM and CRISPR-Cas are encoded by 74.2%^18^ and 39%^19^ of genomes respectively. However, for many systems, the frequency remains to be studied.

One of the reasons for such lack of knowledge can be attributed to the absence of a tool dedicated to the genomic detection of known anti-phage systems. Programs exist for the detection of specific systems such as CRISPR-Cas^20–23^, as well as databases of anti-phage systems such as REBASE for RM^24^ and databases of defense genes such as PADS^25^. However, a single defense gene may not be enough to characterize a functional anti-phage mechanism, therefore it appears more relevant to search for complete systems. The lack of a tool able to find all known anti-phage systems is explained partly by the timeline of the discoveries of the novel systems (mostly since 2018), by their large number (more than 50) and the complex biology of defense systems.

In this study, we developed a tool, DefenseFinder, to detect known anti-phage systems from a genomic sequence. We used this tool, to detect all known anti-phage systems in a database of more than 21 000 complete microbial genomes, describe and analyze their distribution at different phylogenetic scales (from the genome to microbial species and phyla) to provide a systematic and quantitative view of the anti-viral arsenal of prokaryotes.

## Results

### DefenseFinder, a tool to automatically detect known anti-phage systems

We set out to build DefenseFinder, a tool to detect all known anti-phage defense systems in a given genomic sequence. To do so, we used MacSyFinder^26^, a program dedicated to the detection of macromolecular systems. MacSyFinder functions using one model per system. Each model operates in two steps (Figure 1): first the detection of all the proteins involved in a macromolecular system through homology search using HMM profiles; second, a set of decision rules is applied to keep only the HMM hits that satisfy the genetic architecture of the system of interest. This two-steps approach is perfectly adapted for the detection of anti-phage systems, which can exist under different genetic architectures. In fact, it has already been used for the detection of CRISPR-Cas systems^22,26^. We thus built or re-used the HMM profiles of proteins involved in defense systems, and defined specific decision rules for each known anti-phages systems (Figure 1a).

**Figure 1:**
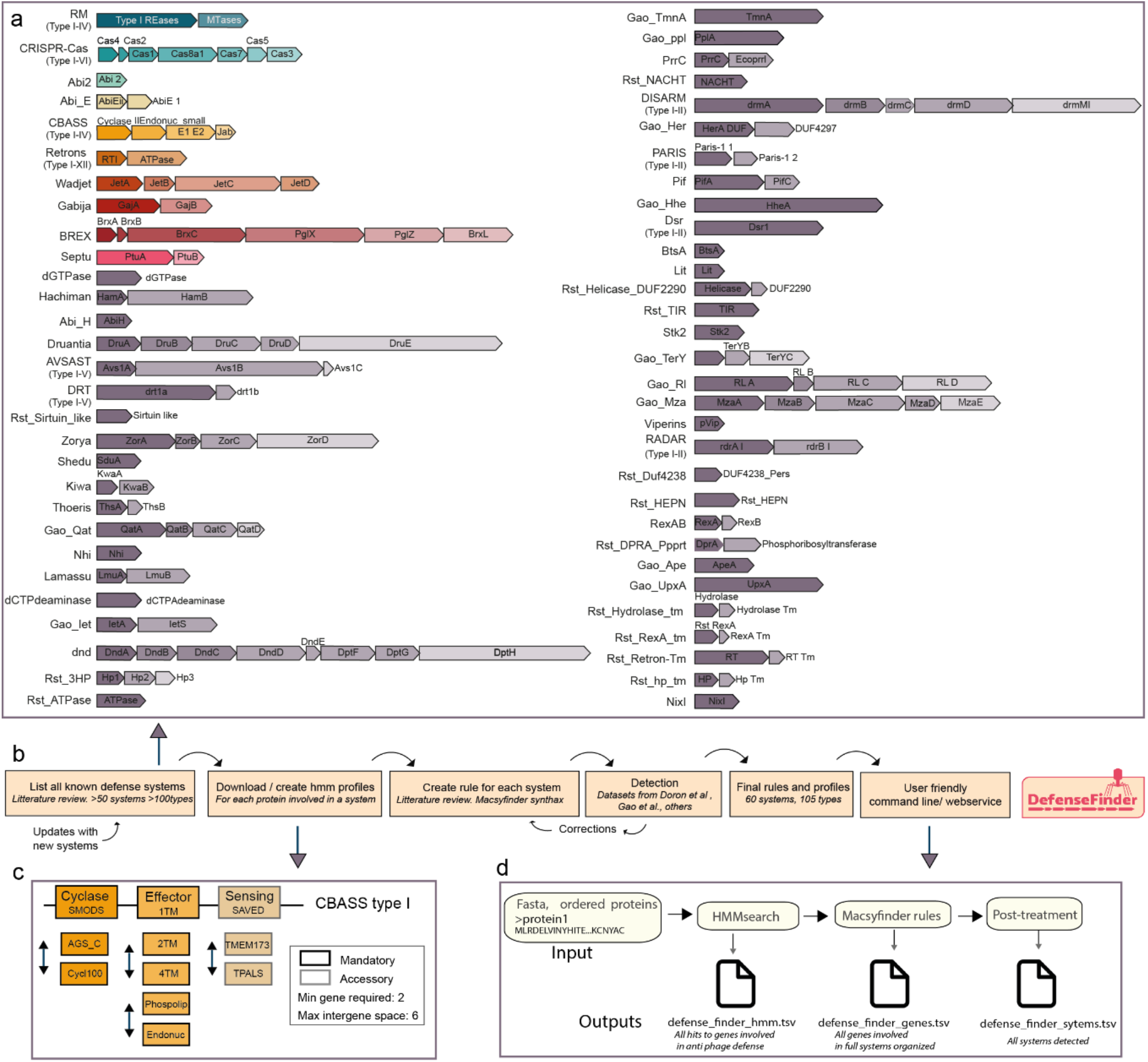
DefenseFinder, a tool to detect all known anti-phage systems. **a.** List of systems included in DefenseFinder. Systems are ordered by frequency in prokaryotic genomes. The 10 most abundant systems are colored (these colors are re-used in Figure4). For systems with several types, each system is represented by one type and other types are indicated in parentheses (Full list, Supplementary Table 1). **b.** Workflow for the creation of DefenseFinder. **c**. Example of a DefenseFinder rule (in the MacsyFinder syntax) for the detection of system CBASS type I. Cyclase and effector proteins are mandatory while the sensing protein is only accessory. This means the system allows for it to be missing in a detected CBASS type I system. Different profiles are recognized for a protein. ex cyclase (SMODS, AGC_C, Cycl100). **d**. DefenseFinder function layout. DefenseFinder takes an ordered multifasta protein file. A search for specific HMM profiles is conducted, the MacSyfinder rules specific for anti-phage systems are applied on the search results, generating three results files.

The building of DefenseFinder required an exhaustive literature search of known anti-phage mechanisms. In this first version of DefenseFinder, we included all described systems discovered before June 2021 (see Material and Methods). We excluded some systems, such as Argonautes and Toxin-Antitoxin, as it is unknown whether all members of such families are involved in anti-phage defense. In total, DefenseFinder detects 60 anti-phage systems (Supplementary Table 1). When available, we also included the types and subtypes of different systems (e.g. CBASS type I, Retron IV, CRISPR-Cas I-E), leading to a total of 105 types of systems (Figure 1b, Supplementary Table 1).

For each of the 105 systems, we defined a model e.g. a set of customized rules and associated protein profiles (see Material and Methods). Briefly, we either used existing pfams/COGs or built custom HMM profiles resulting in a database of 755 profiles. The decision rules are typically defined by a list of mandatory, accessory, or forbidden proteins necessary for the detection of a given system (Figure 1c). For each of the proteins, several homologs can be interchanged. For example, for CBASS type I, two mandatory proteins are necessary (the cyclase and the effector) and a third one is accessory. Each of these proteins have several homologs possible (Effector can be 1TM, 2TM, phospholipase….).

Once a first model is defined, when possible,we evaluate it, against existing datasets^1,19,24,27^ (See Material and Methods). For example, for the 10 systems described in ^1^, where a detailed detection was provided for each system, we downloaded the same database of genomes and used our models to search for these systems. We could then compare our detection and evaluate which systems were for example missing. The model (either profiles or decision rules) could then be adapted. For example, an initial model was missing some occurrences of Shedu system (Supplementary Figure 1). We hypothesized our HMM profile might not have detected some occurrences of SduA (a core protein of Shedu). We built a phylogenetic tree out of all the SduA proteins found in this paper ^1^ and mapped which ones were missed by our detection (Supplementary Figure 1a). We then included additional proteins from under detected clade to our HMM profile. Using similar types of approach, models were improved and we could report a sensitivity for such systems, ranging between 98.7 % (Shedu) and 99,4 % (Shedu) and a high specificity, ranging between 98,5 % (Druantia) and 99,97 % (Thoeris) (Supplementary Figure 1b). Details about validation process for other systems are found in the methods section. All models are under a CC-BY-NC license and available online (https://github.com/mdmparis/defense-finder-models).

The final step was to use such rules and profiles to build as a user-friendly tool to detect anti-phage systems in prokaryotic genomes (Figure 1.d). We provide a command line interface (easily installable through a python package mdmparis-defense-finder) as well as a webservice (https://defense-finder.mdmparis-lab.com/). Both take as an input a protein multifasta file (either one or several genomes at the same time) and generate two types of detection files: a list of detected systems, list of proteins involved in detected systems. The current settings of DefenseFinder are optimized for a conservative detection, as only full systems are present. This can lead to an underdetection of some proteins involved in anti-phage defense. Typically, in defense islands, full systems along single proteins (otherwise involved in an anti-phage defense system) are present. To overcome this, we also propose as an optional output, a list of all the hits to known anti-phage proteins, which can allow a more exhaustive vision of the potential proteins involved in anti-phage defense. The architecture of DefenseFinder is designed for easy and frequent updates of the models, a necessity in the fast-evolving field of anti-phage defense. The webservice will use the most up-to-date rules, and the command line interface can get the most up-to-date rules by calling the option “--update”. The update on the command line is distinct of the update of DefenseFinder allowing to have update of the rules and profile more frequently. Overall, we created DefenseFinder, a program that enables the detection of all known anti-phage systems in prokaryotic genomes.

### The anti-viral arsenal of prokaryotes is highly variable

In order to provide a genome-centric view of the anti-viral arsenal of prokaryotes, we applied DefenseFinder to a database of 21 738 fully sequenced prokaryotic genomes (21 364 bacteria, 374 archaea). We identified 104 737 different defense systems, comprised of 262 960 genes (Supplementary Tables S2, S3). On average prokaryotes encode five anti-phage systems. The number of anti-phage mechanisms per genome varies widely from a minimum of zero (1 567 genomes) to a maximum of 51 in the deltaproteobacteria *Desulfonema limicola*. (Figure 2a, Supplementary Table S4). 75% of prokaryotes encode more than two defense systems.

**Figure 2:**
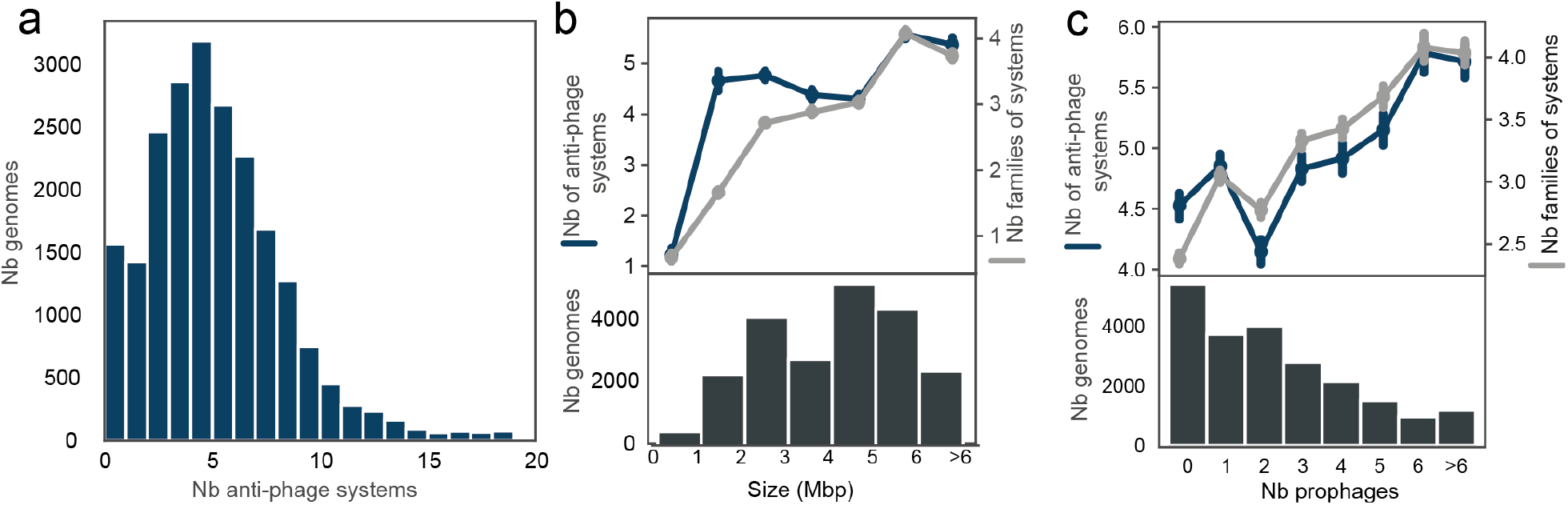
Distribution of anti-phage systems in prokaryotic genomes. **a**. Distribution of the total number of anti-phage systems per genome. The x-axis was cut at 20 for data visualization purposes. Max number is 51 **b**. Average number of anti-phage systems (blue) and number of families of anti-phage systems (grey) are correlated with genome size (Spearman, ρ=0.14 and ρ=0.38, both p-values<0.0001). Bottom plot, distribution of the genome size. **c** Average number of anti-phage systems (blue) and number of families of anti-phage systems (grey) are correlated with number of prophages (spearman, ρ=0.10 and ρ=0.25, both p-values <0.0001). Bottom plot, distribution of the number of prophages in genomes.

To estimate the diversity of this anti-phage arsenal, we computed the number of distinct families of anti-phage systems per genome (e.g. RM, CRISPR-Cas, CBASS). On average, prokaryotes encode three distinct families of anti-phage systems (Supplementary Figure 2). As expected, the number of anti-phage systems correlates positively with the number of families of defense mechanisms (Supplementary Figure 2, Spearman, ρ=0.79 P-value<0.0001). There are some exceptions, with genomes encoding many anti-phage systems but with “low diversity” ie a small number of distinct anti-phage systems families. For example, *Chloroflexus aggregans* DSM 9485 encodes 15 systems but only 3 families of anti-phage systems. This type of genomes typically encodes a wide diversity of RM systems. Indeed, when we focused on the 480 genomes that encode more than 10 anti-phages systems belonging to 4 families or less, we found that for 96% of these genomes, RM systems make up for more than 50% of anti-phage systems. Moreover, 47% of these genomes are from the species *Helicobacter pylori*, which has been described in the past as carrying many RM systems^18^.

We then set out to understand the potential drivers of the number of anti-phage systems in a given genome. The genome size is an important determinant for encoding accessory systems in prokaryotes. It was demonstrated that small genomes encode few CRISPR-Cas and RM^18,19^ and that the number of RM is correlated with genome size^18^. We found this observation can be generalized to the total number of anti-phage systems (Figure 2b blue, Spearman ρ=0.14, p-value<0.0001). However, the size effect is not linear. Very small genomes (<2Mbp) encode very few defense systems, and larger genomes encode more defense systems, but there is a plateau where no size effect is observed for genomes between 2Mbp and 5Mbp. We also observed a stronger positive correlation between the number of families of anti-phage systems and the genome (Figure 2b grey, Spearman ρ=0.38, p-value<0.0001).

We reasoned that the number of anti-phage systems might also be influenced by the diversity of phages a prokaryote might encounter. This can be estimated by focusing on the number of prophages encoded in a given genome. We thus focused on the interplay between anti-phage systems and prophages. To do so, we detected prophages in the genomes of our database using Virsorter2^28^ (see methods). We found 51 582 prophages in 16 315 genomes, with on average two prophages per genome (Supplementary Figure 3) which is in line with previous detection^29^. The number of prophages correlates both with the number of systems (Spearman ρ=0.10, p-value<0.0001) and the number of families (Spearman ρ=0.25, p-value<0.000). We controlled for the effect of the genome size on these parameters using a stepwise forward regression. Both the number of prophages and anti-phage systems are still significantly correlated when taking into account genome size (p-values<0.0001, Figure 2b, Supplementary Figure 3). These results suggest that the anti-viral arsenal of prokaryotes is highly variable and influenced by both genomics traits, such as genome size, and life styles traits, such as the number of prophages present in the genome.

### Families of anti-phage systems have a heterogeneous distribution

We then decided to inspect the distribution of anti-phage systems individually (Figure 3a). RM systems are by far the most abundant as they are present in 80% of prokaryotic genomes, followed by CRISPR-Cas (40%). Apart from these systems, the frequency of the most abundant system drops below 20% (Gabija, Wadjet, Retrons, CBASS, AbiEII, Abi2) ranging from 10% to 17% genomes encoding such systems. The frequencies we find are consistent with what has been described previously^1,6,17–19^. Among the 60 systems detected in this study, 20 are present in less than 1% of the genomes. Such numbers suggest that the diversity of anti-phage system is enormous and many rare systems exist. As RM are the most abundant systems, we checked whether genomes without RM had specific anti-phage mechanisms (Supplementary Figure 4). The most abundant systems in these genomes correspond to the most abundant systems in all prokaryotic genomes suggesting that no specific system replaces RM ones.

**Figure 3:**
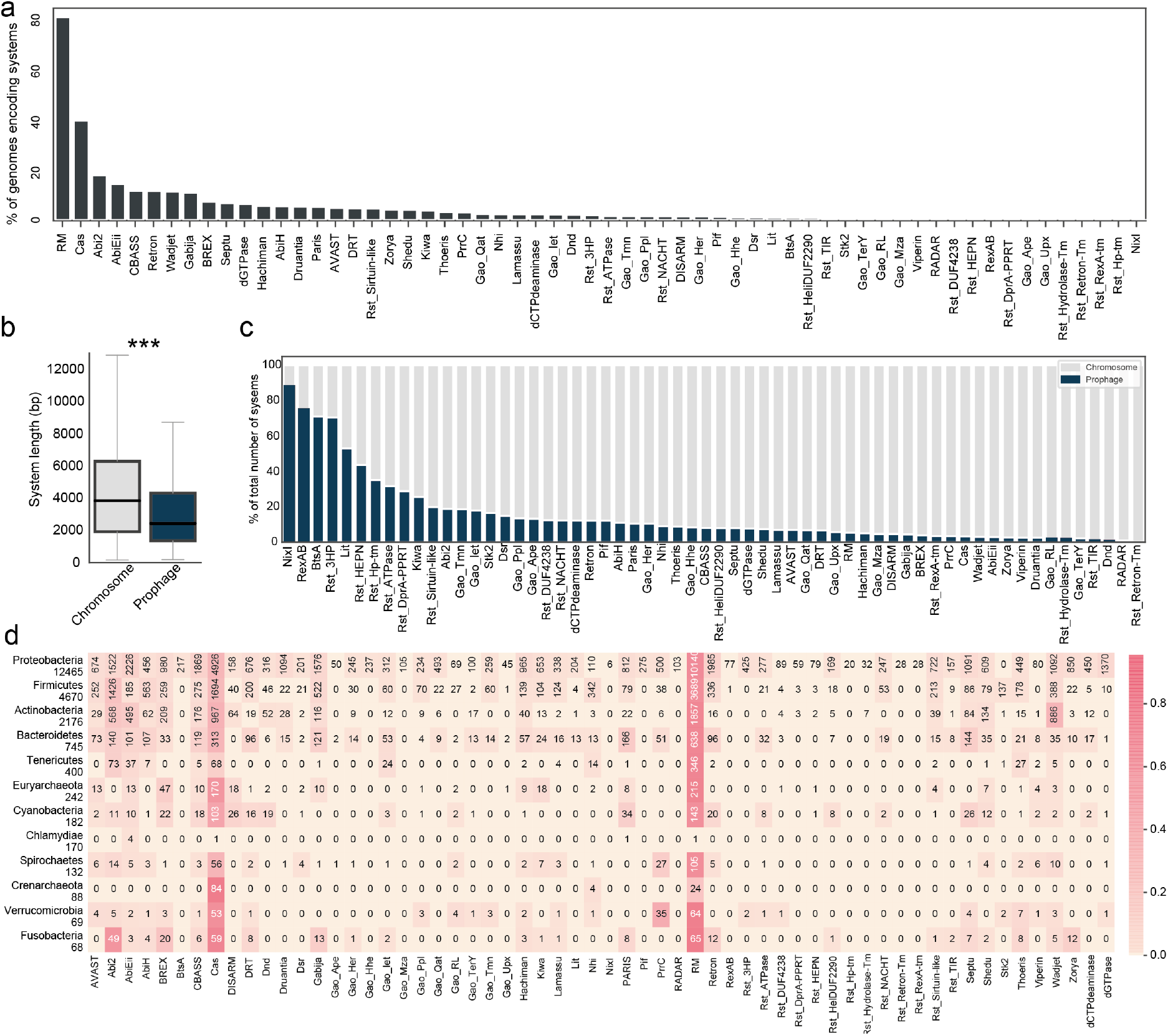
Families of anti-phage systems have an heterogenous distribution. **a**. Frequency of systems in genomes. **b**. System length according to genomic location (***, Wilcoxon p-value<0.0001)). Plasmids were excluded from this analysis. A system was deemed to be part of a prophage when the first and last protein of the system was inside the prophage boundary. The size of a system was computed as the difference between the end of the last protein and the beginning of first protein. **c**. Frequency of genomic location for each system. **d**. Number of systems per prokaryotic phylum. Only phyla with more than 50 genomes are represented. The number of genomes in the dataset is represented under each phylum’s name. The heatmap represents the frequency of each system in a given phylum (“per line”), color legend on the right. Absolute numbers of genomes encoding a given system are indicated in each cell.

Following, the observation that systems’ distribution is contrasted, we set out to understand drivers of such heterogeneity. Several systems recently described (n=17), were discovered in prophages or their parasites^10,12,14,30,31^. These observations could suggest that systems encoded on prophages differ than those encoded in the chromosome. We thus evaluated the anti-phage systems based on their genomic location either prophagic (n=7 530) or chromosomic (n=95 567). It was recently proposed that systems encoded in P2 and P4 (prophages of *E. coli*) could be shorter than other common systems such as CRISPR-Cas^12^. We checked whether this observation could be generalized. Systems located within prophages are shorter (Figure 3.b, median prophagic systems=2 516bp, median chromosomic systems =3 820bp, t-test p-value<0.0001). This is compatible with the size constraints exerted on such elements and suggest that different systems are encoded on the chromosome and prophages. To check this hypothesis, we computed for each system the frequency of the location (either prophagic or chromosomic) (Figure 3.d). For example, out of the 825 Kiwa, 610 are encoded on the chromosome (74%), and 215 in prophages (26%). We found that some systems such as NixI, Rst_3HP, BtsA and RexAB are encoded in majority on prophages, while some systems such as Dnd, viperin or RADAR are almost never found on prophages. The large size of certain systems (Dnd, BREX DISARM) could explain their absence on prophage, but it does not explain why small systems like viperins are not found on prophages in our dataset. It is interesting to notice, that most systems primarily encoded in prophages (NixI, BtsA,RexAB, Lit, Rst system) were discovered in prophages and prophages parasites. While our results are restricted to prophages and not their satellites, they suggest that some systems might be prophage specific.

We then evaluated if phylogeny affects the distribution of anti-phage systems and thus examined the distribution of individual anti-phage systems per phylum (Figure 3c). First, a striking feature is the quasi absence of any anti-phage systems in Chlamydiae. Previous studies also showed a quasi absence of anti-phage systems in genomes of this phyla^17–19,27^. This could be explained by the intracellular lifestyle of such bacteria or by the existence of unknown anti-phage systems for this species. The most widespread systems (RM, CRISPR-Cas) are present in all phyla and are quite abundant in each of these phyla (RM >78% except Chlamydiae and Crenarchaeota; CRISPR-Cas >37% except for Chlamydiae and Tenericutes). Some systems, while less abundant, are widespread across all phyla (such as Abi2, Retrons, CBASS, Hachiman, AVAST…) whereas some other systems are enriched in specific phyla. Typically, many systems such as RexAB, Retron-Tm, Hp-Tm, Ape, are only present in Proteobacteria. This might be explained by a bias of the database towards proteobacterial genomes (n=12 464), which represents 57% of the database; it could indicate the existence of more phylogenetically restricted systems, as most of these systems were discovered in prophages of Proteobacteria; if a system evolves fast it might be missed in distant clades. Finally, some systems seem to be particularly enriched in specific phyla such as BREX in Fusobacteria (Frequency of 29% compared to 8% for all prokaryotes) or Wadjet in Actinobacteria (40% compared to 7% for all prokaryotes). Overall, our results demonstrate that the distribution of anti-phage systems is heterogeneous and influenced by genomic location and phylogeny.

### The anti-viral arsenal of bacteria is species specific

Following our observation that some anti-phage systems are enriched in specific phyla, we decided to focus on the link between anti-phage defense and phylogeny. To do so, we examined the differences between the anti-viral arsenal of diverse species. We selected all the species with more than 100 genomes in our dataset (n=21, see Methods) and established a quantitative comparison between their anti-viral arsenal (Figure 4, Supplementary Figure 5 and 6). Both total numbers of systems and number of different families varies widely between species. For instance, we found no anti-phage system in the 558 genomes of *Bordetella pertussis*, but 16 anti-phage systems on average in *Helicobacter pylori* strains. This is in line with previous reports of *H. pylori* encoding many RM systems^18^. We found species encoding few systems such as *Bacillus subtilis, Staphylococcus aureus, Streptococcus pyogenes* (average number of systems <3) and species encoding many anti phage systems such as *Escherichia coli, Pseudomonas aeruginosa*, or *Neisseria meningitidis* (n>8). Number of systems correlates with diversity of systems within a species, except for three species (*H. pylori, N. meningitidis, C. jejuni*) with many systems (n>6) but only a few families (n<3). Overall these results suggest that species have very diverse types of defense arsenal which could be grouped in three categories: few systems, many diverse systems, many similar systems (Supplementary Figure 6a).

**Figure 4:**
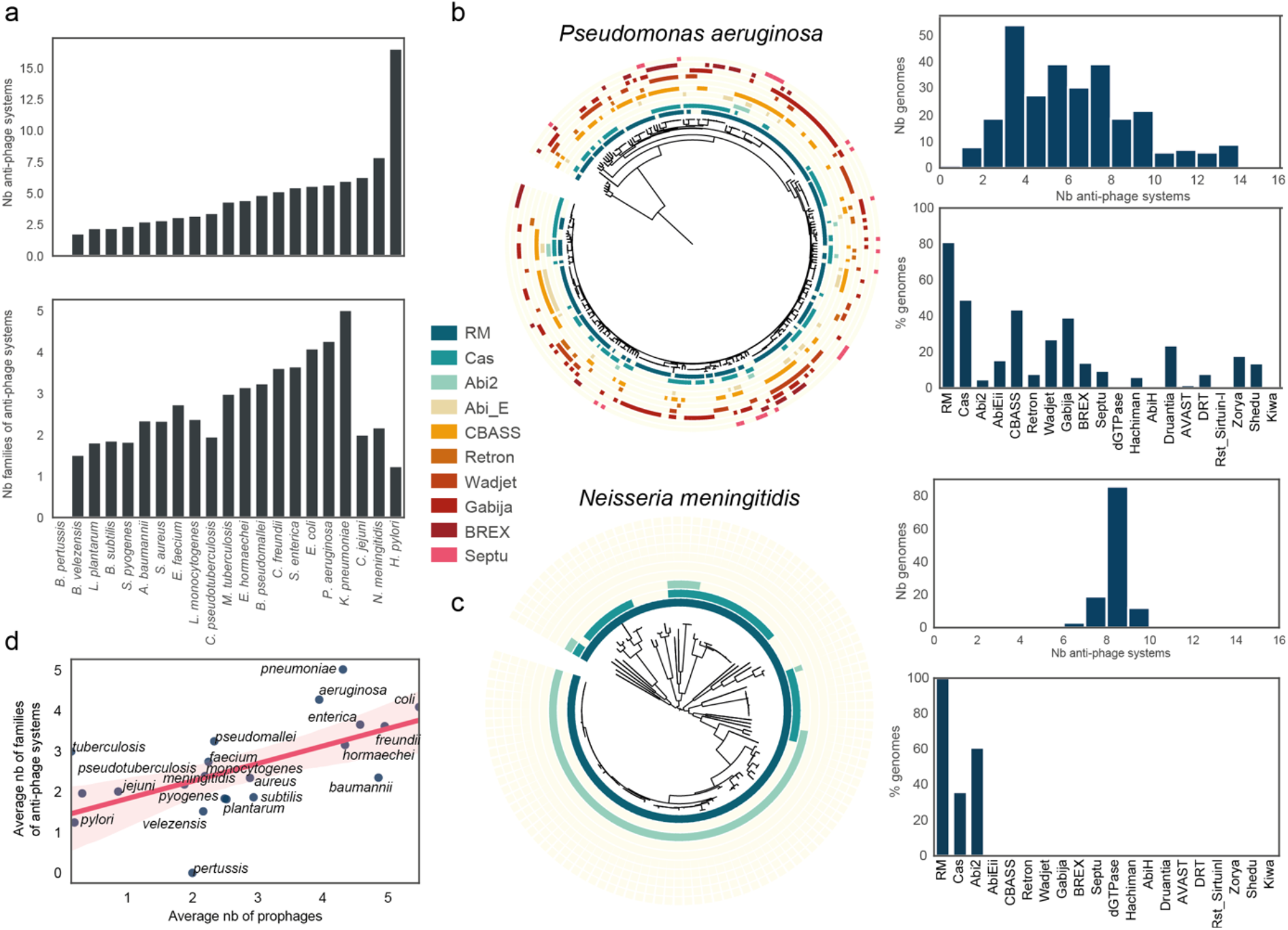
The anti-viral arsenal is species specific. **a**. Number of systems and number of families of systems per species. Species with more than a hundred genomes were selected for this analysis. **b** and **c**. Anti-viral arsenal of two different species: *Pseudomonas aeruginosa* and *Neisseria meningitidis*. Each panel shows the distribution of the total number of systems in the species (top panel), the frequency of the 20 most common anti-phage systems in prokaryotes in this species (bottom panel) and a phylogenetic tree of the species with the presence/absence of the 10 most common anti-phage systems in prokaryotes. **d**. Correlation at the species level between the number of prophages and the families of anti-phage systems (linear regression, pearson r=0.60, p-value=0.002, light pink represent the confidence interval at 95%).

To better understand the different types of anti-viral arsenals, we decided to characterize the anti-phage systems distribution in 15 of these species (See Material and Methods, Figure 3c and d, Supplementary Figures 3 and 4). For each species, we computed 1) the distribution of the total number of systems to evaluate if this was a conserved trait across a species, 2) The frequency of the 20 most common anti-phage systems in prokaryotes in this species 3) A phylogenetic tree based on the core genome of each species where the 10 most common anti-phage systems were reported. These types of representation allow for a comparison between different species. We found very different trends for different species.

An example of the category “many diverse systems” is *P. aeruginosa* (Figure 4c). The number of anti-phage systems varies greatly from one strain to another, going from one to sixteen. Similar to the global distribution, the most common systems are RM and CRISPR-Cas. However, some rarer systems such as CBASS and Gabija are present in more than 40% of the strains while some common systems such as dGTPase are absent from these genomes. Our phylogenetic analysis reveals a patchy distribution of anti-phages systems even in closely related strains, suggesting high rate of horizontal gene transfer. For example, the two closely related strains *P. aeruginosa* Ocean-1155 (GCF_002237405.1) and *P. aeruginosa* C79 (GCF_007833895.1) have a very different anti-viral arsenal (respectively 5 systems with 1 RM, 1 Abi2, 1 AbiEii, and a Retron type V vs 10 systems including BREX, Gabija, Shedu, ATPase and Iet). This is in line with previous observations that *P. aeruginosa* closely related strains could have diverse anti-viral arsenal^32^, but demonstrates that this trend concerns the entire species not a specific part of it. To control that our results were not due to biases in the number or diversity of genomes, we analyzed the correlation at the species level between the diversity in the anti-viral arsenal and the phylogenetic distance (Supplementary Figure 6b). To do so, we computed the Bray-Curtis distance of all pairs of anti-viral arsenals as well as the phylogenetic distance between all strains (Supplementary Figure 4). We found no correlation between these values for *P. aeruginosa*. We found a similar trend for other species such as *E. faecium* and *A. baumanii* (Supplementary Figure 6b).

The study of the anti-viral arsenal of *N. meningitidis* uncovered very different conclusions. The number of anti-phage systems is almost constant with a very narrow distribution centered around 8 systems. We found only three families of anti-phage systems, RM, CRISPR-Cas and Abi2, the most common systems in prokaryotes. Contrary to *P. aeruginosa*, the presence of anti-phage systems is very dependent on the phylogeny (Supplementary Figure 6, spearman, ρ=0.68, p-value<0.0001). For example, 63 phylogenetically closely related strains have exactly 5 RM type II, 2 RM Type III, and 1 Abi2. We found a similar trend for species including *C. jejuni, S. pyogenes* and *S. aureus* (Supplementary Figure 6b).

We then set out to understand what could drive such different strategies to fight phage infection. We observed that for all prokaryotes, genome size and number of prophages influence the number of anti-phage systems and of families, we postulated that these factors could influence the anti-viral arsenal of species. At the species level, the total number of anti-phage systems does not correlate with the number of prophages or the size of genomes (Supplementary Figure 5). However, the number of families of anti-phage systems correlates with both genome size and number of prophages (Figure 7e, Supplementary Figure 5, spearman, ρ=0.66, ρ=0.64, p-values =0.001 and 0.002). This suggests that the diversification of the anti-phage arsenal, not the number of anti-phage systems, is impacted by these factors.

## Discussion

We created a tool to detect all known anti-phage systems. To our knowledge, this represents the first tool to perform the detection for all known anti-phage systems. We hope that the availability of this tool in command line (for big data analysis) and through a web-service (for occasional detection) will allow the community to learn more about specific strains and bring new insights on large datasets. To that end, DefenseFinder is quite adapted as it runs in less than a day on 10 000 genomes on a regular laptop (using 4 CPU). A major challenge for such program is to adequately reflect the state of the literature in the field. Because the literature on this particular topic is ever changing, we took advantage of the architecture proposed by MacSyFinder to facilitate regular updates.

Many features of most anti-phage systems are still not understood such as their diversity or molecular mechanisms. This lack of knowledge leads to challenges in the detection, as some might exist under unknown forms and would not be detected by our program. It is also possible that several systems will end up being regrouped in bigger families of anti-phage systems. All our models are openly available, and we hope the community will propose novel and better models or point out to missing or inaccurate ones. Another hurdle of our program is the heavy reliance on computer inference, as we sometimes have only one or a couple systems that were experimentally validated. Similarly, for many systems, the accuracy of the detection could not be properly evaluated as no existing ground truth was available, which only experimental work can provide. Finally, the systems are only detected through proteins and no nucleic acid components is detected here (such as msDNA for retrons or CRISPR arrays for CRISPR-Cas). This will be added in a future version.

We detected and analyzed thousands anti-phage systems in fully sequenced bacterial and archaeal genomes. We used the entire RefSeq complete genome database, which is known to be biased towards specific prokaryotes, notably to over-represent cultivable bacteria. However, it represents the first census of the full anti-viral arsenal of prokaryotes. While we know this census will change in its details as many anti-phage systems probably remain to be discovered, we expect that the general trends that were observed in our study will not change much. Indeed, the most recent systems to be discovered are present in less than 15% of the genomes. Thus, the discovery of additional systems, might impact deeply the phage-host interactions for a specific strain or species, but the general numbers for all prokaryotes should not change drastically. Metrics we used in our study are also subjects to limitations. We evaluated the diversity of the anti-viral arsenal using families of anti-phage systems which we define as ensemble of systems with similar molecular mechanisms (RM, CRISPR, CBASS). However, much diversity notably in terms of molecular mechanism exists within such families and could be inspected. Second, we used the number of prophages as an estimate of phage diversity. It has been shown that is some cases, species have few prophages but many virulent phages^33^, this metrics does not take this aspect into account.

Despite these limitations, our data provides a quantitative description of the anti-viral arsenal of prokaryotes which will serve to answer fundamental questions in anti-phage biology. It confirms that only a few anti-phage systems are very abundant. If many rare systems exist, it is highly possible that many more remains to be discovered. We showed that several anti-phage systems are enriched in prophages or in specific phyla. Understanding the causes of this enrichment might reveal specific evolutionary constraints imposed on this system. Another observation from this census is the diversity of anti-phage arsenals at the species level. While specific enrichment in some anti-phage systems had been shown for specific anti-phage systems such as RM or CRISPR-Cas^19^, our results suggest specific mechanisms are acting at the level of this anti-viral arsenal.

We previously postulated the existence of a potential “Pan immune system”^32^ where “*the ‘effective’ immune system is not the one encoded by the genome of a single microorganism but rather by its pan-genome, comprising the sum of all immune systems available for a microorganism to horizontally acquire and use*.”. Several recent studies reported diversity and dynamics of anti-phage systems in natural isolates of *Vibrio*^13,34,35^ compatible with such hypothesis. Our current study suggests that this hypothesis could be relevant only for a subset of species, while other evolutionary strategies might be at play for others. Our results suggest that a high diversity of phages could lead to a diversification of the anti-viral arsenal. Other studies will be needed to shed light on such evolutionary dynamics which could have important implications for phage therapy.

Overall, our study provides both a tool and a census for the detection of anti-phages systems paving the way to quantitative examination of several additional hypotheses in the field of anti-phage defense, such as the phenomenon of defense islands or co-occurrences between anti-phage systems. Our tool will allow new genomic insights in this booming field.

## Materials and methods

### Choice of anti-phage systems

We chose to include all anti-phage systems described with at least one experimental evidence of the anti-phage function (before June 2021). As the field is fast-evolving, we decided to also include systems described in preprints. Some systems such as Argonautes and Toxin-Antitoxin have not been included yet, as there is still some controversy if all members of such families are involved in anti-phage defense. In total, DefenseFinder detects 60 families of anti-phage systems (Supplementary Table 1). While our aim is to be exhaustive, it is possible that some systems were missed, we call on the community to help us complete and correct the list.

### Building DefenseFinder HMM Models

The protein profiles used were either retrieved from existing databases (PFAM^36^, COG^37^) or built from scratch when no adequate profiles existed (see below for details on the building of HMM profiles and Supplementary Table 1).

New protein profiles for the proteins involved in anti-phage systems were built using a homogeneous procedure. We collected a set of sequences from the protein family that were representative of the diversity of the bacterial taxonomy. Homologous proteins were aligned using MAFFT v7.475^38^ (default options, mode auto) and then used to produce protein profiles with Hmmbuild (default options) from the HMMer suite v3.3^39^. To ensure a better detection we curated each profile manually by assigning a GA score (used with the hmmsearch option --cut_ga). GA score defines the threshold above which a hit is considered significant. This threshold was determined manually by inspecting the distribution of the scores.

Protein scrapping was done using different methods depending on the system knowledge (Details in Supplementary Table 1). For systems from^1^, dGTPase^8^, dCTPdeaminase^8^, BREX^3^, part of Cyclic-oligonucleotide-based anti-phagesignalling systems (CBASS)^17^, all the reverse transcriptases of retrons, BtsA^10^, viperins^7^ and DISARM^2^, we used a subset (between 20 and 100 proteins) of the proteins available in the supplementary data of each publication. We then tested if the HMM allows for detection of all known occurrences of such proteins. If a lot of proteins were undetected, we added proteins reported in the supplementary materials but not detected through our HMM to the list of sequences for the alignment and subsequent HMM generation.

For AbiEii, AbiH,Abi2, Stk2, Pif, Lit, PrrC, RexAB, part of CBASS, part brxA, and brxB from BREX, PARIS (AAA15 and AAA21), we used PFAM available at (http://pfam.xfam.org/) or the sequence available on COG (https://www.ncbi.nlm.nih.gov/research/cog-project/). For part of BREX, DndABCDEFGH^40^, we searched for proteins with this name available on NCBI and curated manually such list. For systems when only one sequence was provided such as Gao’s systems^4^, Rousset’s systems^12^, Dnd type SspBCDE, part of retrons, the protein sequence was BLASTed. Between 20 and 50 sequences with high coverage were selected. For retrons other than reverse transcriptase, we used the IMG genome neighborhood feature to get adjacent protein of the reverse transcriptase and repeated the BLAST process. For CAS systems, HMM protein profiles were downloaded from^22,41^. All hmm profiles used are available at https://github.com/mdmparis/defense-finder-models.

### Building DefenseFinder rules of detections

We defined genetic organization rules based on the literature (Supplementary table S1). MacSyfinder allows for two types of genetic components, “mandatory” and “accessory”. Given the wide diversity of genetic organization of anti-phage systems, initial rules were written differently for major types of systems. Typically, for small systems (less than 3 proteins), the number of mandatory proteins required were strict whereas for bigger system (such as Druantia), the number of proteins required did not always required all components to be present. For CAS systems, all models previously defined in CasFinder v2.0.2 include in CRISPRCasFinder^22^ have been rewritten to be compatible with the new version of MacSyfinder v2.0rc4^26^ and updated to take into account the most recently proposed nomenclature^41^. As a result, this new version CasFinder v3.0.0 (used in DefenseFinder) allows to detect 6 different types (I-VI) and 33 different subtypes. All DefenseFinder rules used are available at https://github.com/mdmparis/defense-finder-models.

### Validation of DefenseFinder models

Following an initial design of rule for each system, we ran an initial detection and evaluated the results. When available, this initial detection was compared to other existing datasets.

For example, specificity and sensitivity were evaluated for each system from ^1^ and is reported in Supplementary Figure 1. Sensitivity was defined as the percentage of system detected in ^1^ and by DefenseFinder. Specificity was defined as the ratio between the number of genomes where a system was detected by DefenseFinder in genomes where in ^1^ had not detected any and all the genomes where ^1^ has not detected the system. For CRISPR-Cas systems, detection was compared to CasFinder v2 from ^22^. For RM systems, we ran DefenseFinder on REBASE^24^ (Supplementary Figure 1c). Overall, we report a 91% sensibility (319466 systems detected out of 349327). Our profiles are underdetecting some distant forms of RM type IV (23% of the RM type IV on REBASE are missed) however, when such sequences were added to the profile we observed an important drop in specificity which led us to the current trade-off.

Another type of verification was to check if we could accurately type systems for which different subtypes are available. For example, retrons are a family of defense systems recently described^6,27^. A classification in twelve types was published earlier this year accompanied by the detection of types in thousands of fully sequenced genomes generating a first estimate of the frequency of such types. We applied DefenseFinder on RefSeq, and evaluated the frequency of such types of retrons. The comparison between these two results revealed almost no differences in frequencies of retrons types (Supplementary Figure 1). Some subsystems are less detected by DefenseFinder such as type X or type XIII. However, the database used for detection is not the same for the two analysis and the phylogenetic repartition could modify the type distribution.

When no other datasets was available, the quality of the detection was estimated by checking different factors such as the number of occurrences found, size of proteins and systems. Each rule was thus refined through trial and error cycles to reach a final stable version.

## Availability

DefenseFinder online is available at https://defense-finder.mdmparis-lab.com/

DefenseFinder command line is available through *pip install mdmparis-defense-finder*

DefenseFinder documentation is available at https://github.com/mdmparis/defense-finder

DefenseFinder models are available at https://github.com/mdmparis/defense-finder-models.

## Data

We analyzed 21 738 complete genomes retrieved in May 2021 from NCBI RefSeq representing 21 364 and 374 species of Bacteria and Archaea (http://ftp.ncbi.nih.gov/genomes/refseq/bacteria/).

## Detection of anti-phage systems

We used DefenseFinder v 0.0.11 (models v0.0.3, August 2021) to search for anti-phage systems in the RefSeq database. To do so, we first formatted this database under a gembase format (see DefenseFinder documentation). We then ran DefenseFinder with the –db-type gembase. We provide the results of this detection in Supplementary Tables 2-5.

## Phylogenetic analysis

We used PanACoTA^42^ version 1.2.0 to build phylogenies for 15 bacterial species (*Escherichia coli, Pseudomonas aeruginosa, Streptococcus pyogenes, Salmonella enterica, Listeria monocytogenes, Helicobacter pylori, Mycobacterium tuberculosis, Neisseria meningitidis, Staphylococcus aureus, Bacillus subtilis, Campylobacter jejuni, Klebsiella pneumoniae, Bacillus velezensis, Acinetobacter baumannii, Enterococcus faecium*). PanACoTA allows building phylogenetic trees using core genomes. For each of the species, we took all genomes under a nucleic acid format in NCBI (fna) and annotated them using prodigal (PanACoTA annotate options --cutn 10000 --l90 400 –prodigal). We then computed the pangenome and coregenome (PanACoTA pangenome; PanACoTA corepers; with default parameters). Finally, we aligned the coregenome (PanACoTA align, default parameters) and computed a phylogenetic tree (PanACoTA tree, -b 1000). For this step PanACoTA, uses IQTree^43^, (version 2.1.4) and the following option (iqtree -m GTR -bb1000 -st DNA)

## Detection of prophages

Putative prophages were detected using VirSorter v2.2.2^28^. Results were filtered to exclude the least confident predictions (we kept max score > 0.8, size <200kb) which may be prophage remnants or erroneous assignments.

## Figures and statistical analysis

Figures were made with matplotlib v3.3.2^44^ and seaborn v0.11.0^45^. Data analysis and statistics analysis were done using pandas 1.1.3^46^ and scipy 1.5.2^47^. Phylogenetic trees were plotted using ITOL^48^. Statistical significance were adjusted by Bonferonni correction.

## Ackowledgements

We would like to thank Eduardo PC Rocha, Gal Ofir and members of MDM lab for fruitful discussion and review of the manuscript; Nitzan Tal for very fruitful advice on data visualization and the manuscript; Adrien Bernheim for the logo design of DefenseFinder, Aurélien Hervé for the help on the webserver interface and Bertrand Neron and Sophie Abby for the development, maintenance and advice on MacSyFinder.

## References

1. Doron, S. et al. Systematic discovery of antiphage defense systems in the microbial pangenome. Science 359, eaar4120 (2018).

2. Ofir, G. et al. DISARM is a widespread bacterial defence system with broad anti-phage activities. Nat. Microbiol. 1–9 (2017). doi:10.1038/s41564-017-0051-0

3. Goldfarb, T. et al. BREX is a novel phage resistance system widespread in microbial genomes. EMBO J. 34, 169–183 (2015).

4. Gao, L. et al. Diverse enzymatic activities mediate antiviral immunity in prokaryotes. Science 369, 1077–1084 (2020).

5. Cohen, D. et al. Cyclic GMP–AMP signalling protects bacteria against viral infection. Nature 574, 691–695 (2019).

6. Millman, A. et al. Bacterial retrons function in anti-phage defense. Cell 1–11 (2020). doi:10.1101/2020.06.21.156273

7. Bernheim, A. et al. Prokaryotic viperins produce diverse antiviral molecules. Nature 589, 120–124 (2021).

8. Tal, N. et al. Antiviral defense via nucleotide depletion in bacteria. bioRxiv (2021).

9. Severin, G. et al. A Broadly Conserved Deoxycytidine Deaminase Protects Bacteria from Phage Infection. bioRxiv (2021).

10. Owen, S. V. et al. Prophage-encoded phage defence proteins with cognate self-immunity. bioRxiv (2020). doi:10.1101/2020.07.13.199331

11. Bari, S. M. N. et al. A unique mode of nucleic acid immunity performed by a single multifunctional enzyme. bioRxiv (2020).

12. Rousset, F., Dowding, J., Bernheim, A., Rocha, E. P. C. & Bikard, D. Prophage-encoded hotspots of bacterial immune systems. bioRxiv 2021.01.21.427644 (2021). doi:10.1101/2021.01.21.427644

13. LeGault, K. N. et al. Temporal shifts in antibiotic resistance elements govern phage-pathogen conflicts. Science 373, eabg2166 (2021).

14. LeGault, K. N., Barth, Z. K., DePaola IV, P. & Seed, K. D. A phage parasite deploys a nicking nuclease effector to inhibit replication of its viral host. bioRxiv (2021). doi:: https://doi.org/10.1101/2021.07.12.452122

15. Ofir, G. et al. Antiviral activity of bacterial TIR domains via signaling molecules that trigger cell death. bioRxiv 2021.01.06.425286 (2021).

16. Kronheim, S. et al. A chemical defence against phage infection. Nature 564, 283–286 (2018).

17. Millman, A., Melamed, S., Amitai, G. & Sorek, R. Diversity and classification of cyclic-oligonucleotide-based anti-phage signalling systems. Nat. Microbiol. 5, 1608–1615 (2020).

18. Oliveira, P. H., Touchon, M. & Rocha, E. P. C. The interplay of restriction-modification systems with mobile genetic elements and their prokaryotic hosts. Nucleic Acids Res. 42, 10618–10631 (2014).

19. Bernheim, A., Bikard, D., Touchon, M. & Rocha, E. P. C. Atypical organizations and epistatic interactions of CRISPRs and cas clusters in genomes and their mobile gemetic elements. Nucleic Acids Res. 1–13 (2019). doi:10.1093/nar/gkz1091

20. Russel, J., Pinilla-Redondo, R., Mayo-Muñoz, D., Shah, S. A. & Sørensen, S. J. CRISPRCasTyper: Automated Identification, Annotation, and Classification of CRISPR-Cas Loci. Cris. J. 3, 462–469 (2020).

21. Mitrofanov, A. et al. CRISPRidentify: Identification of CRISPR arrays using machine learning approach. Nucleic Acids Res. 49, (2021).

22. Couvin, D. et al. CRISPRCasFinder, an update of CRISRFinder, includes a portable version, enhanced performance and integrates search for Cas proteins. Nucleic Acids Res. 46, W246–W251 (2018).

23. Padilha, V. A., Alkhnbashi, O. S., Shah, S. A., De Carvalho, A. C. P. L. F. & Backofen, R. CRISPRcasIdentifier: Machine learning for accurate identification and classification of CRISPR-Cas systems. Gigascience 9, 1–12 (2020).

24. Roberts, R. J., Vincze, T., Posfai, J. & Macelis, D. REBASE-a database for DNA restriction and modification: Enzymes, genes and genomes. Nucleic Acids Res. 43, D298–D299 (2015).

25. Zhang, Y. et al. PADS Arsenal: a database of prokaryotic defense systems related genes. Nucleic Acids Res. 48, D590–D598 (2020).

26. Abby, S. S., Néron, B., Ménager, H., Touchon, M. & Rocha, E. P. C. MacSyFinder: A Program to Mine Genomes for Molecular Systems with an Application to CRISPR-Cas Systems. PLoS One 9, e110726 (2014).

27. Mestre, M. R., González-Delgado, A., Gutiérrez-Rus, L. I., Martínez-Abarca, F. & Toro, N. Systematic prediction of genes functionally associated with bacterial retrons and classification of the encoded tripartite systems. Nucleic Acids Res. 48, 12632–12647 (2020).

28. Guo, J. et al. VirSorter2: a multi-classifier, expert-guided approach to detect diverse DNA and RNA viruses. 1–13 (2021).

29. Moura de Sousa, J. A., Pfeifer, E., Touchon, M. & Rocha, E. P. C. Causes and Consequences of Bacteriophage Diversification via Genetic Exchanges across Lifestyles and Bacterial Taxa. Mol. Biol. Evol. 38, 2497–2512 (2021).

30. Parma, D. H. et al. The Rex system of bacteriophage λ: Tolerance and altruistic cell death. Genes Dev. 6, 497–510 (1992).

31. Uzan, M. & Miller, E. S. Post-transcriptional control by bacteriophage T4: MRNA decay and inhibition of translation initiation. Virol. J. 7, 1–22 (2010).

32. Bernheim, A. & Sorek, R. The pan-immune system of bacteria: antiviral defence as a community resource. Nat. Rev. Microbiol. 18, 113–119 (2020).

33. Touchon, M., Bernheim, A. & Rocha, E. P. Genetic and life-history traits associated with the distribution of prophages in bacteria. ISME J. 10, 2744–2754 (2016).

34. Hussain, F. A. et al. Rapid evolutionary turnover of mobile genetic elements drives microbial resistance to viruses. bioRxiv 1–18 (2021).

35. Piel, D. et al. Genetic determinism of phage-bacteria coevolution in natural populations. bioRxiv (2021). doi:: https://doi.org/10.1101/2021.05.05.442762;

36. Finn, R. D. et al. Pfam: The protein families database. Nucleic Acids Res. 42, 222–230 (2014).

37. Galperin, M. Y., Makarova, K. S., Wolf, Y. I. & Koonin, E. V. Expanded Microbial genome coverage and improved protein family annotation in the COG database. Nucleic Acids Res. 43, D261–D269 (2015).

38. Katoh, K., Misawa, K., Kuma, K. & Miyata, T. MAFFT: a novel method for rapid multiple sequence alignment based on fast Fourier transform. Nucleic Acids Res. 30, 3059–3066 (2002).

39. Eddy, S. R. Accelerated profile HMM searches. PLoS Comput. Biol. 7, (2011).

40. Xiong, X. et al. SspABCD–SspE is a phosphorothioation-sensing bacterial defence system with broad anti-phage activities. Nat. Microbiol. 5, 917–928 (2020).

41. Makarova, K. S. et al. Evolutionary classification of CRISPR–Cas systems: a burst of class 2 and derived variants. Nat. Rev. Microbiol. 18, 67–83 (2020).

42. Perrin, A. & Rocha, E. P. C. PanACoTA: a modular tool for massive microbial comparative genomics. NAR genomics Bioinforma. 3, nqaa106 (2021).

43. Minh, B. Q. et al. IQ-TREE 2: New Models and Efficient Methods for Phylogenetic Inference in the Genomic Era. Mol. Biol. Evol. 37, 1530–1534 (2020).

44. Hunter, J. D. Matplotlib: A 2D Graphics Environment. Comput. Sci. Eng. 9, 90–95 (2007).

45. Waskom, M. et al. mwaskom/seaborn: v0.8.1 (September 2017). (2017). doi:10.5281/zenodo.883859

46. McKinney, W. {D}ata {S}tructures for {S}tatistical {C}omputing in {P}ython. in {P}roceedings of the 9th {P}ython in {S}cience {C}onference (eds. van der Walt, S. & Millman, J.) 56–61 (2010). doi:10.25080/Majora-92bf1922-00a

47. Virtanen, P. et al. {SciPy} 1.0: Fundamental Algorithms for Scientific Computing in Python. Nat. Methods 17, 261–272 (2020).

48. Letunic, I. & Bork, P. Interactive tree of life (iTOL) v3: an online tool for the display and annotation of phylogenetic and other trees. Nucleic Acids Res. 44, W242–W245 (2016).

