## Supplementary materials for "Systematic and quantitative view of the antiviral arsenal of prokaryotes"

### Supplementary material

##### Supplementary Table 1
List of systems, rules and HMM profiles used in DefenseFinder

##### Supplementary Table 2
Anti-phage systems detected.

##### Supplementary Table 3
Genes involved in anti-phage systems

##### Supplementary Table 4
Anti-phage systems per genome

##### Supplementary Table 5
Detection of prophages

### Supplementary Figures

##### Supplementary figure 1: Validation of DefenseFinder models
**a.** Phylogenetic tree of SduA. Dark blue was found by an initial round of the DefenseFinder models, light blue was missed (compared to detection from Doron et al) **b.** Table of sensitivity and sensibility for systems from Doron et al. **c.** Validation of the detection of RM **d.** Comparison of distribution of retron subtypes.

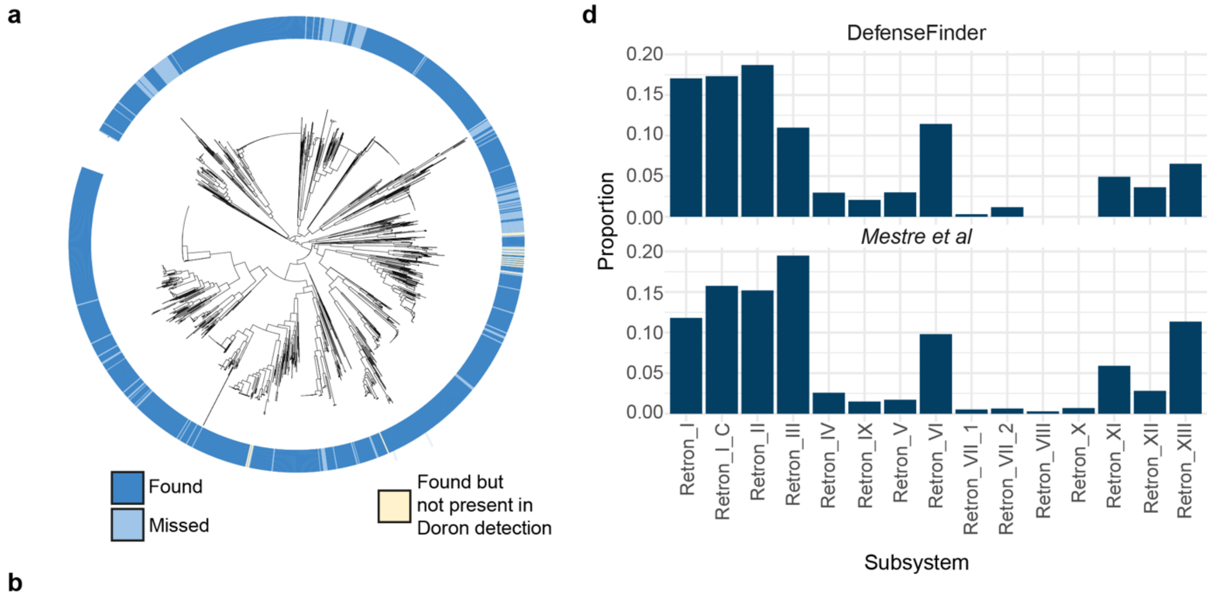


**b**

| System | Druantia | Gabija | Hachiman | Kiwa | Lamassu | Septu | Shedu | Thoeris | Wadjet | Zorya |
|---|---|---|---|---|---|---|---|---|---|---|
| Sensitivity | 99,0 % | 99,1 % | 98,8 % | 98,9 % | 99,2 % | 99,4 % | 98,7 % | 98,5 % | 99,3 % | 99,0 % |
| Specificity | 98,5 % | 99,2 % | 99,9 % | 99,5 % | 99,9 % | 99,3 % | 99,9 % | 99,9 % | 99,8 % | 99,9 % |

**c**

| R-M Protein | Type I M | Type I R | Type II M | Type II R | Type III M | Type III R | Type IV |
|---|---|---|---|---|---|---|---|
| Sensitivity | 98,3 % | 98,9 % | 91,1 % | 89,1 % | 97,6 % | 83,7 % | 77,4 % |

**Supplementary figure 2: Families of anti-phage systems are correlated with the total number of anti-phage systems**

**a.** Distribution of the number of families anti-phage systems per genome. The x-axis was cut at 20 for data visualization purposes. **b.** Correlation between the families of anti-phage systems and the total number of anti-phage systems ( Spearman ρ=0.79 P-value<0.0001).

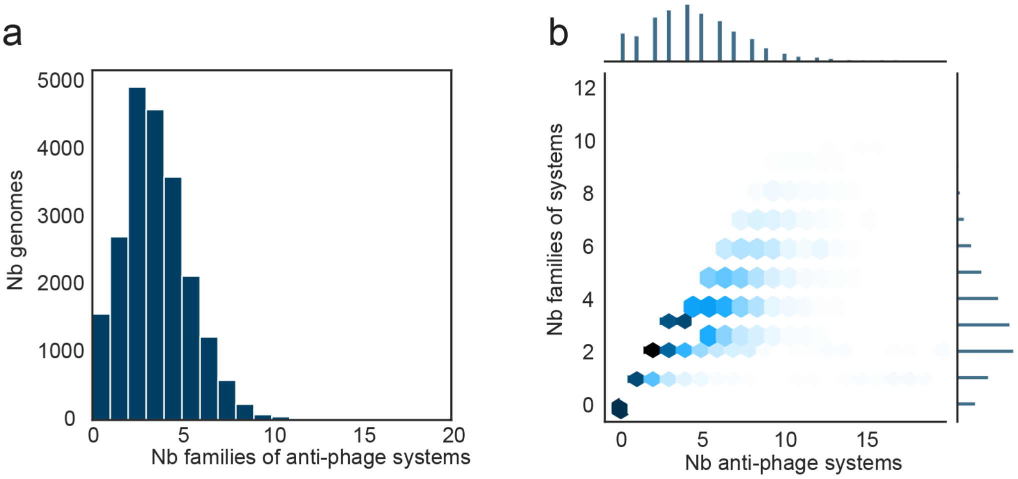


**Supplementary figure 3: Prophage detection**

**a.** Distribution of the number of prophages per genome. **b.** Correlation between number of prophages and genome size.

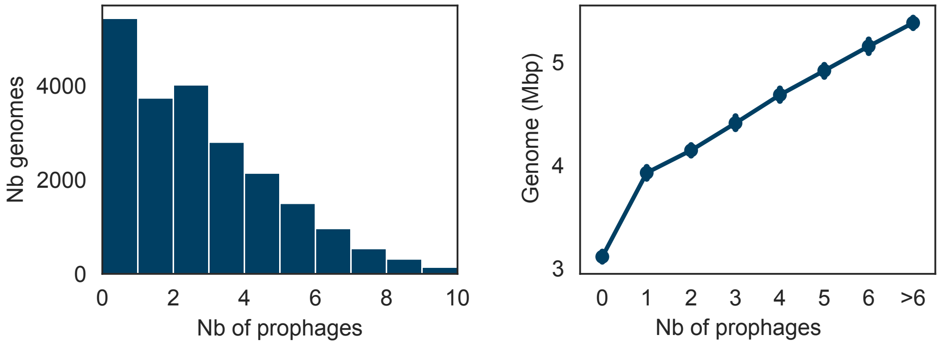

### Supplementary figure 4: Distribution of anti-phage systems in genomes without RM

**a.** Distribution of anti-viral systems. **b.** Distribution per phyla

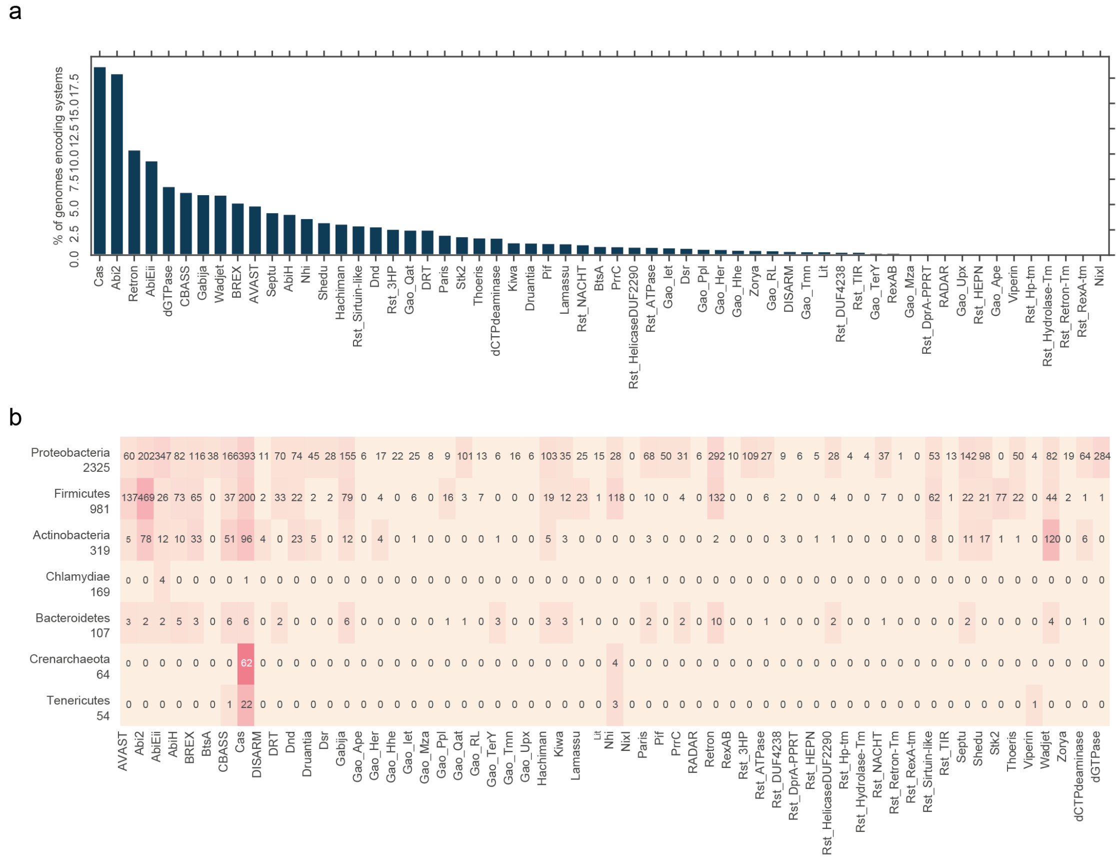

#### Supplementary figure 5: Anti-viral arsenal of diverse bacterial species

Each panel shows the distribution of the total number of systems in the species (top panel), the frequency of the 20 most common anti-phage systems in prokaryotes in this species (bottom panel) and a phylogenetic tree of the species with the presence/absence of the 10 most common anti-phage systems in prokaryotes.

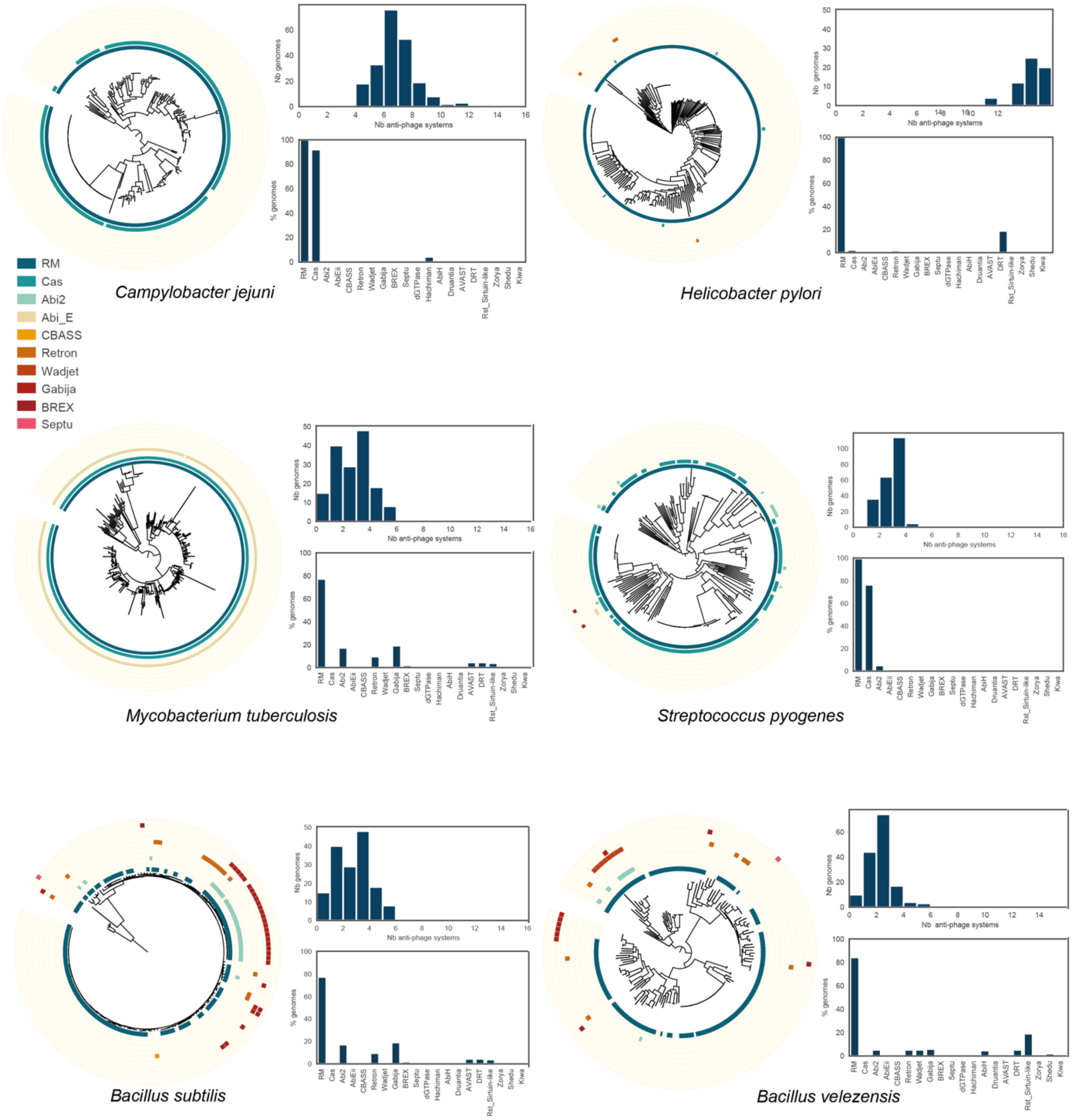

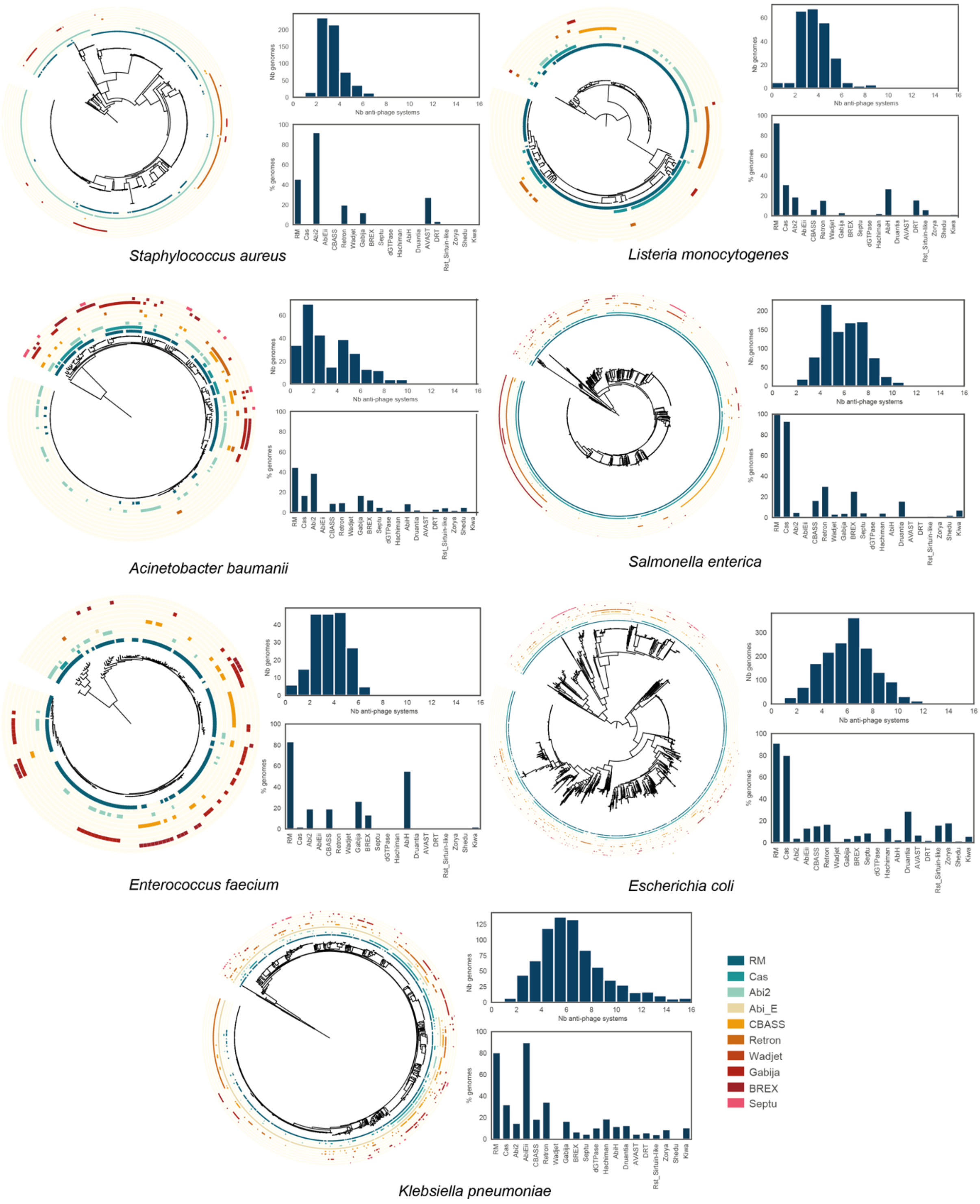

Nb genomes
Nb anti-phage systems
% genomes
RM
Cas
Abi2
AbiEii
CBASS
Retron
Wadjet
Gabija
BREX
Septu
dGTPase
Hachiman
AbiH
Druantia
AVAST
DRT
Rst_Sirtuin-like
Zorya
Shedu
Kiwa
Staphylococcus aureus
Listeria monocytogenes
Acinetobacter baumanii
Salmonella enterica
Enterococcus faecium
Escherichia coli
Klebsiella pneumoniae
RM
Cas
Abi2
Abi_E
CBASS
Retron
Wadjet
Gabija
BREX
Septu

**Supplementary Figure 6: Anti-viral arsenal of bacterial species are diverse**

**a.** Scatter plot for the anti-viral arsenal of prokaryotes. **b.** Correlation between the phylogenetic distance and the Bray-Curtis distance of the anti-viral arsenal of diverse species. Each plot corresponds to one species. For each species, the Bray-Curtis distance (dist_BC) of all pairs of anti-viral arsenals was computed as well as the phylogenetic distance (dist_phylo) between all strains. Each line corresponds to a pearson fit of the data.

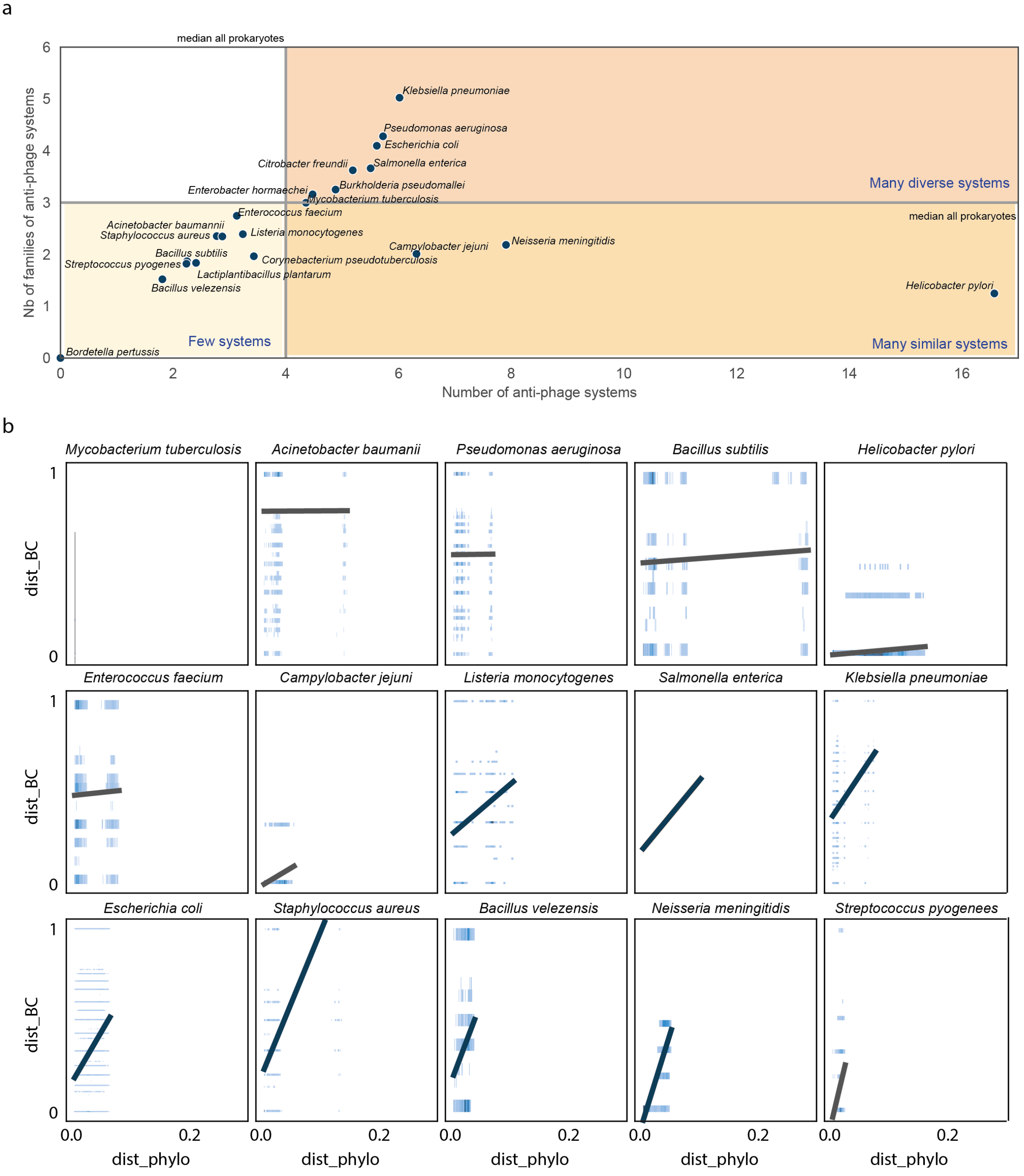

**Supplementary figure 7: Determinants of the anti-viral arsenal of bacterial species**

**a.** Genome size of bacterial species. **b.** Correlation between the families of anti-phage systems and the genome size (Linear regression pearson r=0.65, p-value=0.0014). **c.** Correlation between the number of anti-phage systems and the number of prophages (linear regression r=-0.2 p-value=0.4). **d.** Correlation between the number of anti-phage systems and the genome size (linear regression r=-0.16 p-value=0.5).

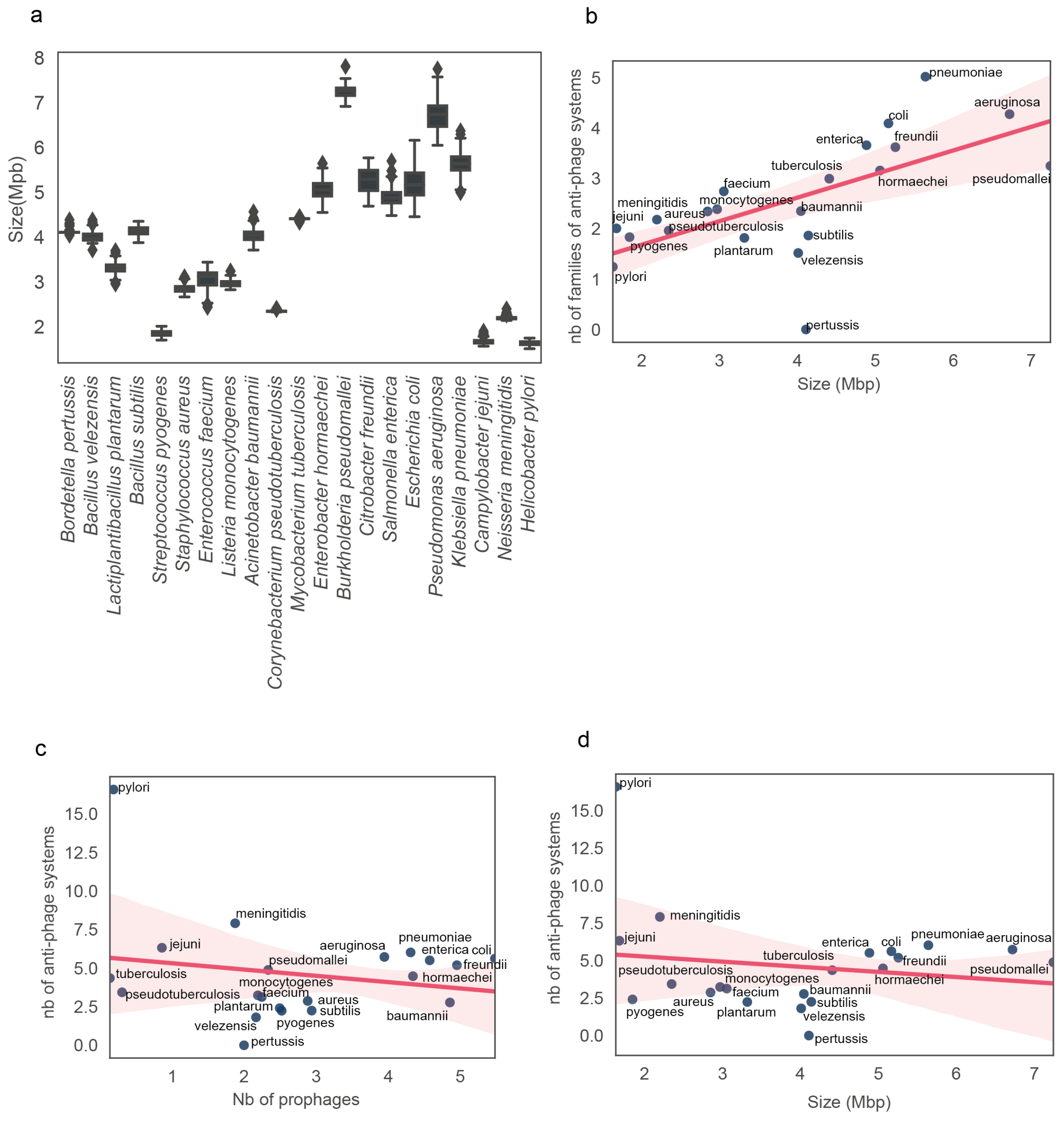
